# Context effects in pitch discrimination reflect response bias not sensory bias

**DOI:** 10.64898/2026.07.02.735981

**Authors:** Coral E. Dirks, Daniel R. Guest, Andrew J. Oxenham

## Abstract

Context effects are ubiquitous across sensory systems and reflect a general encoding principle for both simple and complex stimuli. One simple context effect, contraction bias, manifests in two-interval perception tasks as a bias of the perceived magnitude of the first stimulus toward the center of the overall magnitude range. The underlying cause of contraction bias is unclear. One explanation is that a listener’s magnitude estimate of the first stimulus is combined with a perceptual anchor, usually the mean stimulus magnitude, biasing it toward the anchor (sensory model). An alternative explanation is that a listener’s response criterion shifts, based on the magnitude of the stimulus pair, relative to the mean magnitude of the stimuli range (decision model). Two pitch-discrimination experiments were performed to test these hypotheses in the auditory domain. The first was a forced-choice discrimination task, where listeners were asked to identify the higher or lower tone in a pair. The second was a same-different task where listeners indicated whether or not the two tones in a pair differed in frequency. Contraction bias was observed in the higher-lower discrimination task, even after extensive perceptual training with feedback. In contrast, no contraction bias was observed in the same-different task. Computational models of the sensory and decision hypotheses were fit to data from both experiments. The sensory model captured the pattern of results the higher-lower experiment but erroneously predicted a contraction bias in the same-different task. The decision model produced similar predictions to the sensory model in the higher-lower task but correctly predicted no contraction bias in the same-different task, and produced lower prediction errors and more stable parameter estimates in both paradigms. Overall, the results suggest that the underlying nature of the contraction bias may reflect decision, rather than sensory, biases based on the context.

**AUTHOR SUMMARY:** Every day, people combine experiential knowledge of the world and incoming sensory information to assess their environment. Interestingly, the environment itself can substantially alter how incoming sensory information is processed and perceived. These so-called context effects occur in all sensory modalities. Here, we examine one such context effect, known as “contraction bias”, in a simple pitch-discrimination task. Prior work argued that the contraction bias occurs because observers make biased estimates of stimulus magnitudes due to the influence of recent stimulus history. By combining behavioral data and computational modeling, we offer a different explanation, more consistent with the available data, which posits that the bias can be accounted for by systematic shifts in observers’ response strategies with changes in relative stimulus magnitudes. The results shed light onto how humans compare and process sensory features within their environment.

## INTRODUCTION

Humans are remarkably sensitive to changes in the frequency of a sound. When presented with two successive tones, trained listeners can tell whether the frequency increased or decreased even with changes as small as 0.5% [1, 2]. This difference is an order of magnitude smaller than a semitone or half-step (∼6%) – the smallest interval used in Western music. Despite this impressive accuracy, our discrimination abilities can be strongly affected by the context in which the tones are presented. For instance, when the two tones are presented in a higher frequency range than preceding tones, the first tone is more often reported as lower in frequency than the second; conversely, when the two tones are presented in a lower frequency range than the preceding tones, then the first tone is more often heard as higher than the second tone. This effect has been termed contraction bias [3] and belongs to a family of context effects known collectively as time-order errors, which have been studied across many sensory modalities for over a century in dimensions such as size, weight, and frequency [4].

Contraction bias is particularly pronounced when the two stimuli are separated in time by at least several seconds [5, 6]. A common explanation has been that the recent (and rapidly decaying) sensory memory of the first tone is combined with the “expected” frequency, based on the aggregate frequencies of the preceding tones, i.e., the center of the range of preceding frequencies, so that the perceived frequency of the second tone in the pair is compared not with the direct sensory representation of the first tone, but with a weighted combination of the first tone and a “perceptual anchor” based on prior exposure [4, 7]. This long-standing concept has been formalized within the more modern framework of Bayesian inference [8].

In contrast to the idea that a perceptual anchor requires extended exposure to become established, Raviv et al. [3] found evidence for contraction bias in as few as 160 trials, or less than five minutes, and with the two tones separated by less than 1 second. It is unlikely that listeners had enough experience to create stable predictions about how the stimuli behaved; therefore, Raviv et al. proposed an alternative hypothesis: rather than pooling information from all prior observations, listeners may rely on a perceptual anchor that is formed from a weighted average of the frequencies presented in the past few trials, with stimuli in more recent history being weighted more heavily. This hypothesis was supported in a follow-up study that also used frequency discrimination as the model [9].

Although their data and model were broadly consistent with the literature, some aspects of the Raviv et al. [3] data are puzzling. In particular, their data showed substantial contraction bias, despite a relatively short gap of less than a second between the two stimuli in each trial. In addition, their participants had average discrimination thresholds of around 13%, which is more than a whole tone (or two semitones) on the diatonic scale, and thus more than an order of magnitude poorer than thresholds obtained by practiced participants [2], even when tones are roved in frequency between trials [10–12].

Since the participants were not trained and the experiment itself was very short, it is possible that listeners did not fully understand the task. For instance, rather than identifying the higher tone, participants may simply have judged whether the entire tone pair was above or below the mean of the frequency range. Such a strategy could explain the direction of the observed bias and could also explain the surprisingly poor discrimination exhibited by the participants in Raviv et al.’s [3] study.

The first aim of the current study was to determine whether the large bias effects observed by Raviv et al. persist once participants are better trained and exhibit more sensitivity to small frequency differences. The second aim was to attempt to dissociate sensory bias (based on perception) from response bias (based on decision criteria). Our results show that significant bias can be measured in listeners with high sensitivity to frequency differences, even after perceptual training, and that the bias cannot be explained simply by listeners basing their judgments on the average frequency of the tone pair, relative to previous pairs. Moreover, the bias was only observed in an up-down task, and not in a same-different task, suggesting the presence of response, rather than sensory, bias. An alternative model derived from signal detection theory and consistent with previous work [13] is proposed to account for the results from the different experiments presented here.

## RESULTS

### JUST NOTICEABLE DIFFERENCES IN FREQUENCY

In the first experiment, listeners were presented with two tones, separated by a silent gap. In one task, listeners were asked to select the higher of the two tones. In the other task, listeners were asked to select the lower of the two tones. These two tasks were used to test whether listeners were biased towards selecting the right button if the tone pair was higher than preceding tones and selecting the left button if the tone pair was lower than preceding tones, in line with the mapping on a piano keyboard. Such a bias would produce what appears to be a contraction bias for the more common “select the higher tone” task, but would produce the opposite bias in the novel “select the lower tone” task. We refer to both as “up-down” tasks, as listeners had to discriminate whether the frequency contour was rising (Low-High tone order) or falling (High-Low tone order). The two tasks (“select higher” and “select lower”) were tested in counterbalanced order across participants. This experiment consisted of four test sessions. In all cases, the average or center frequency (CF) of the two tones ranged between 800 and 1200 Hz, with the CF selected at random from ten possible values on each trial. For the first and second test sessions, the frequency difference between the two tones (Δ*f*) was fixed within a block of trials at 1, 2, or 5% of the CF. In the third test session, Δ*f* values of 0.5, 0.75, and 1.5% were tested. In the fourth and final test session, Δ*f* values of 0, 0.75, and 1.5% were tested.

Just-noticeable differences (JNDs) were estimated for each test session by fitting a psychometric function to the listeners’ data, pooled across tasks and across CF values. Estimated psychometric functions appear as solid curves in Figure 1. The Δ*f* at which performance was predicted to be 70.7% correct was defined as the JND, because this is the level of performance tracked by a widely used 2-down 1-up adaptive procedure for estimating threshold [14] and corresponds roughly to a value of 0.77 for the index of sensitivity, *d’* [15]. The estimated JNDs were 1.00%, 0.87%, 0.72%, and 0.58% for the first, second, third, and fourth test sessions, respectively, and appear as dashed vertical lines in Figure 1. These results show that the listeners improved their performance on average over the four test sessions, but were already highly sensitive to changes in frequency even in the first session, with thresholds more than an order of magnitude better than those reported by Raviv et al. [3].

**FIGURE 1.**
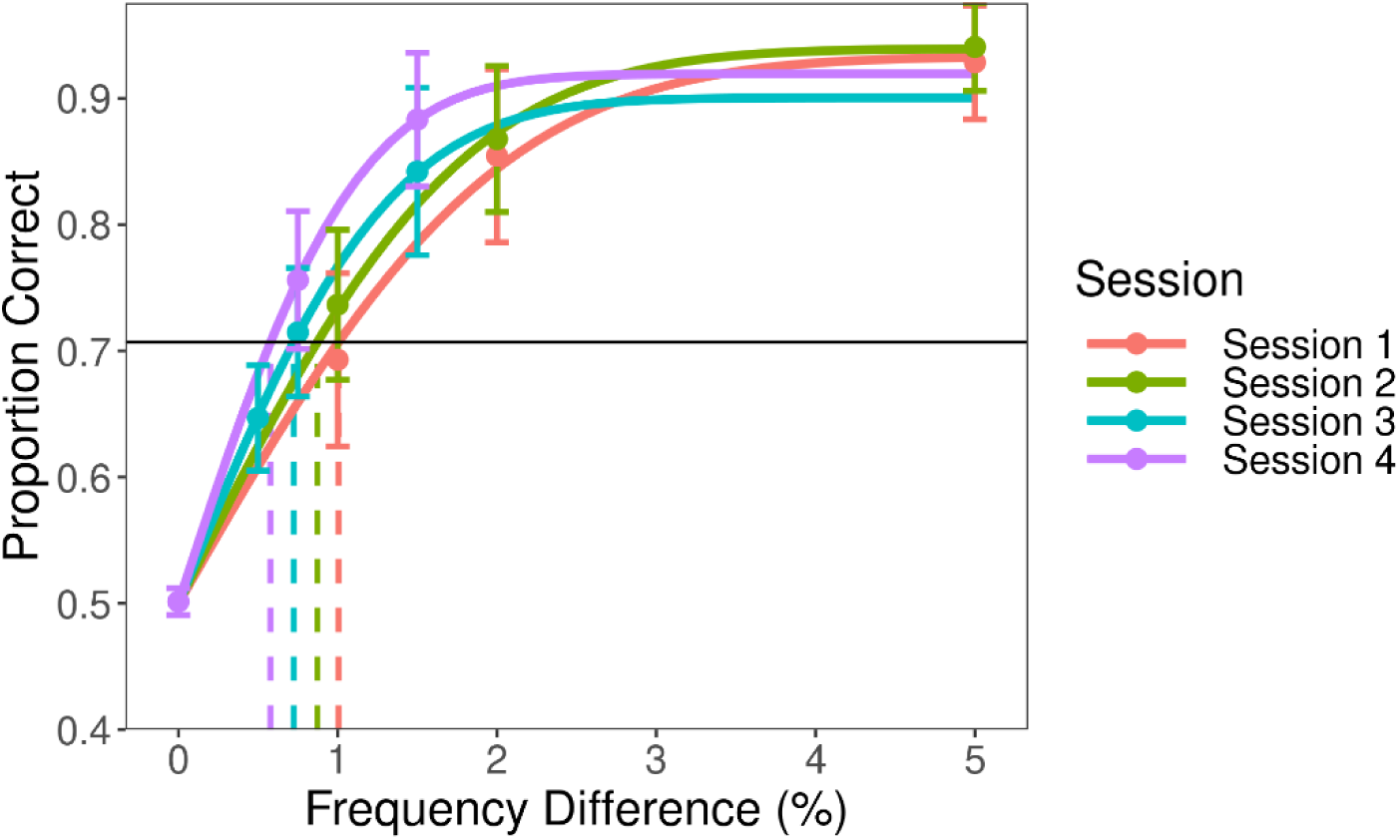
Proportion of correct responses in the frequency-discrimination task as a function of frequency difference (*Δf*). Mean performance appears as solid dots. Data are separated according to test session by color and error bars represent ±1 standard error of the mean. Individual data were used to estimate JND according to Dai and Micheyl [16]; model fits to the group-average percent-correct values appear as curves. The JND was defined as the value of *Δf* at which performance was equal to 70.7% correct (horizontal black line). The vertical dashed lines show the JNDs for the curves fitted to the mean data from each session.

A linear mixed-effects model was applied to the individual log-transformed JND estimates, with fixed effects of test session and task (i.e., whether the listener was instructed to select the higher or lower tone in a given trial) as well as random intercepts for each listener. Results revealed a significant main effect of test session [*F*(3, 119.37) = 3.39, *p* = 0.020] but no significant main effect of task [*F*(1, 119.00) = 3.84, *p* = 0.052] and no significant interaction between task and test session [*F*(3, 119.00) = 0.75, *p* = 0.52]. *Post hoc* contrast tests revealed that JNDs improved significantly from the first to fourth test session [estimated ratio = 0.72, *F*(1, 119.54) = 10.07, *p* = 0.012], although no pairwise contrasts between the other test sessions were significant. These results confirm that the listeners had excellent frequency discrimination that improved significantly with practice between the first and last sessions.

### BIAS REVEALED WITH UP-DOWN EXPERIMENT

Data from individual trials in the up-down experiment were separated by trial order (High-Low vs. Low-High), task (select high or low tone), session number, and tone-pair CF, and then analyzed to answer three specific questions. First, did the contraction bias appear in listeners with small JNDs? The average JND in the Raviv et al. [3] study was greater than 13% and listeners showed a very large bias, with performance near ceiling in conditions that favored the bias (e.g., rising tone pairs in the upper portion of the frequency range) and near chance in unfavorable conditions (e.g., falling tone pairs in the upper portion of the frequency range). The sensitivity of our listeners was an order of magnitude better; good pitch discrimination implies an accurate sensory representation that could reduce the influence of any nonsensory bias. Second, was the bias affected by the instructions given to listeners? If the contraction bias is a perceptual phenomenon, listeners should continue to indicate that the tone in the first interval is closer to the mean magnitude of the frequency range irrespective of whether they are tasked with indicating the interval containing the higher tone or the interval containing the lower tone. Conversely, if the bias reflects an erroneous mapping, or task confusion, on the part of the listeners, the bias may be reversed in sign by changing the task instructions. Third, does contraction bias remain after extensive perceptual training? It is possible that the magnitude of the bias might decrease with increasing training with feedback and improved perceptual sensitivity.

Data from the up-down frequency-discrimination experiment appear again in Figure 2, but this time they are broken down by the CF of the tone pair and by the tone order (i.e., the order in which the tones were presented, either Low-High or High-Low). The effects of bias are illustrated in Figure 2: the proportion of correct responses to low-high trials increases with increasing CF of the tone pair (dashed lines), while the proportion correct to high-low trials decreases with increasing CF (solid lines). To determine the statistical significance of the effects of the various experimental parameters, an analysis of deviance was performed using a generalized linear mixed-effects model (GLMM; see Methods for details) that was fit to all the data from the up-down discrimination experiment. The analysis of deviance revealed significant main effects of Δ*f* (χ_6_^2^= 7702.01, *p* < 0.001), tone-pair type (low-high or high-low) (χ_1_^2^ = 17.71, *p* < 0.001) and task (select higher or lower tone) (χ_1_^2^ = 20.59, *p* < 0.001). It also revealed significant two-way interactions between Δ*f* and tone-pair type (χ_1_^2^= 25.09, *pp* = 0.0030), tone-pair type and task (χ_6_^2^ = 133.86, *p* < 0.001), and tone-pair type and CF (χ_1_^2^ = 19.75, *p* < 0.001), a significant three-way interaction between Δ*f*, tone-pair type, and CF (χ_6_^2^ = 60.18, *p* < 0.001), and a significant four-way interaction between all terms in the model (χ_6_^2^= 21.96, *p* = 0.010). No other model terms reached significance.

**FIGURE 2.**
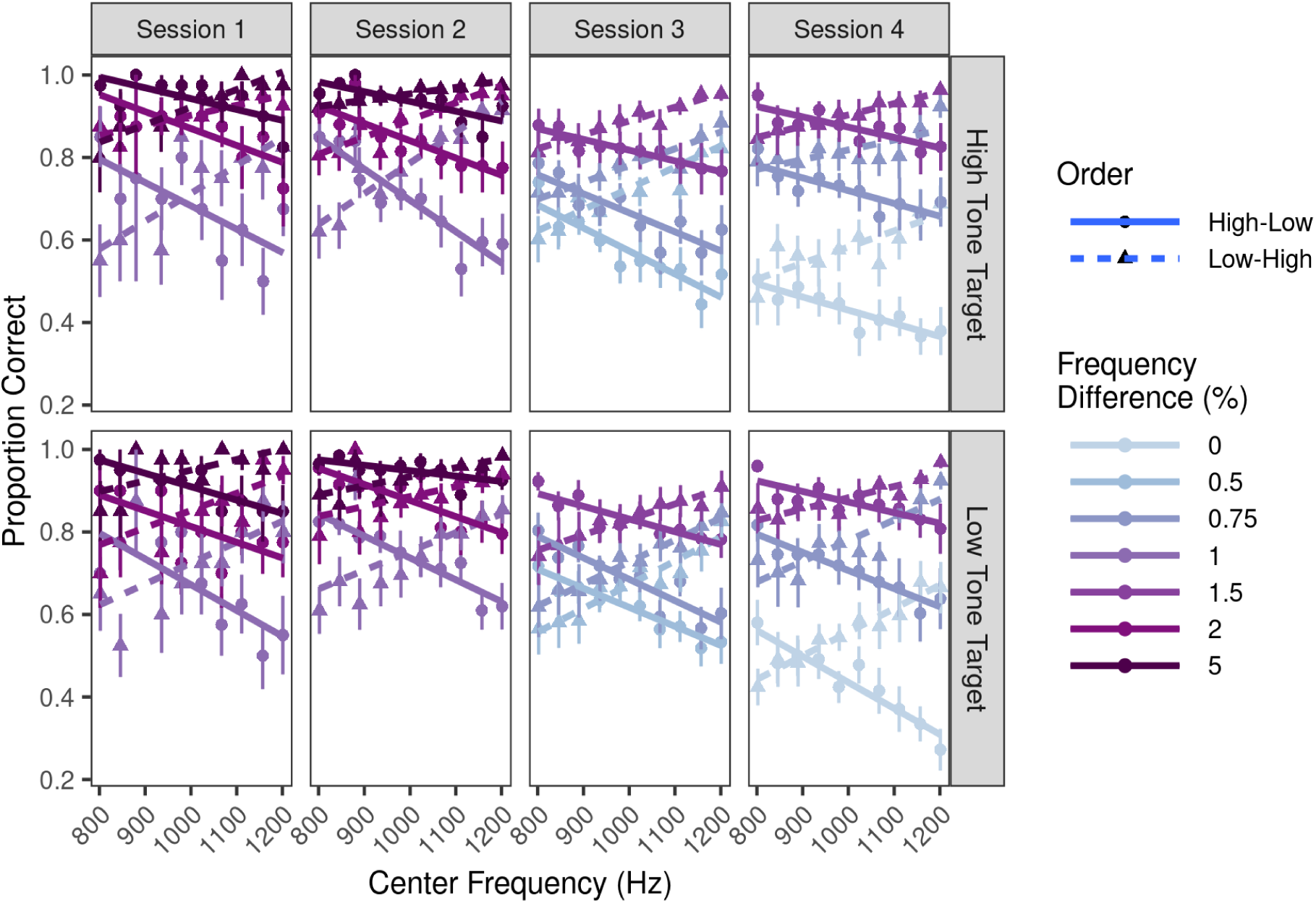
Proportion of correct responses as a function of the tone pair center frequency in the up-down frequency-discrimination experiment. Data are separated into columns according to test session (1-4) and rows according to task (select high or low tone in each pair). Color denotes the frequency difference between two tones on any given trial, with more saturated colors indicated greater values of Δ*f*. Lines are linear fits to the data, and line type differentiates the direction of the tone-pair pitch contour, with solid and dashed lines corresponding to trials where the higher-frequency tone was in the first and second interval, respectively. Error bars represent ±1 standard error of the mean across the 20 (Sessions 1-3) or 14 (Session 4) participants.

*Post hoc* tests were performed to investigate the three key hypothesized features of the data. First, is the contraction bias reported in Raviv et al. [3] present in our data? In other words, did the probability of a correct response increase with increasing CF in Low-High trials and decrease in High-Low trials? Contrast tests confirmed the impression from Figure 2 that such bias is present in our data. The slope parameter associated with CF (hereafter CF slope), which was centered at 1000 Hz (see Methods for details), was significantly different from zero and positive in the Low-High condition (χ_1_^2^ = 27.10, *pp* < 0.001), indicating that increases in CF were associated with increases in the probability of a correct response in the Low-High trials.

Likewise, the CF slope was significantly different from zero and negative in the High-Low condition (χ_1_^2^ = 18.13, *p* < 0.001), indicating that increases in CF were associated with *decreases* in the probability of a correct response in the High-Low trials. An interaction contrast test confirmed that CF slopes in the Low-High and High-Low condition were significantly different from each other (χ_1_^2^ = 23.07, *p* < 0.001).

Second, does the sign of the bias depend on the task? Contrast tests revealed no significant differences in the CF slope parameter for task in either High-Low trials (χ_1_^2^ = 0.64, *p* = 0.85) or in Low-High trials (χ_1_^2^= 0.12, *p* = 0.85). This result suggests that listeners were similarly biased by the CF of the tone pair whether they were tasked with indicating the interval containing the lower tone or the interval containing the higher tone. This finding rules out the possibility that the contraction bias observed here and previously was due to listeners’ confusion about the task or response bias, as described above, with the right button being associated with higher tones pairs.

Third, is the bias robust, even in listeners with high sensitivity to frequency differences, and does it persist after with training and feedback? To address this question, we first fitted sub-models to each session separately (resulting in four total sub-models) and then tested the significance of the bias in each sub-model to confirm that the bias was present in each session of testing. We confirmed that the CF slopes were significant and in the expected direction for the first (High-Low,χ_1_^2^= 10.57, *p* = 0.0034; Low-High, χ_1_^2^ = 28.01, *p* < 0.001), second (High-Low, χ_1_^2^ = 14.64, *p* < 0.001; Low-High, χ_1_^2^ = 28.98, *p* < 0.001), third (High-Low, χ_1_^2^ = 11.65, *p* = 0.0019; Low-High, χ_1_^2^= 20.13, *p* < 0.001), and fourth sessions (High-Low, χ_1_^2^= 5.78, *p* = 0.048; Low-High, χ_1_^2^= 6.89, *p* = 0.043). Second, we fitted two additional sub-models, one including only the data collected during the first and second sessions (as these two sessions of the experiment tested the same Δ*f* values) and another including only the data collected during the third and fourth sessions for the Δ*f* values of 0.75% and 1.5% (as these two sessions of the experiment both tested those Δ*f* values). We tested whether the bias (i.e., CF slope in each tone pair type) differed significantly between test sessions. These tests revealed no significant differences in CF slope from the first session to the second session for the High-Low trials (χ_1_^2^ = 0.39, *p* = 1.00) or for the Low-High trials (χ_1_^2^ = 1.17, *p* = 1.00). Likewise, for the High-Low trials no significant change in CF slope was detected from the third session to the fourth session (χ_1_^2^ = 4.10, *p* = 0.13); however, in the Low-High trials a significant (although small) decrease in the CF slope was observed (χ_1_^2^ = 8.04, *p* = 0.018). Collectively, these results suggest that although practice may decrease the size of the bias effect to some extent, the bias largely persists even after extensive experience with the task.

It is interesting to note that the largest contraction bias was observed in the fourth session at Δ*f* = 0%. In these trials, the tones in each interval were the same frequency, yet the computer software randomly designated one interval as the “target” interval. As expected, overall performance was at chance (50% correct), but performance diverged strongly between the two tone orders as a function of CF, highlighting the strong effect of the bias. In other words, even when the two tones presented to listeners had the same frequency, listeners tended to report the first tone as lower than the second tone if the tones were above the mean of the frequency range and the first tone as higher than the second tone if the tones were below the mean of the frequency range.

Overall, the results reveal several important new findings about the well-known contraction bias effect. First, contraction bias persists even when sensitivity to frequency differences is high (i.e., JNDs less than 1%). Second, the participant instructions (“select the higher tone” or “select the lower tone”) had no significant effect on the direction of the bias, suggesting that the observed bias did not simply reflect a mapping between particular buttons and the overall pitch height of the tone pair. Finally, the bias persisted even after 4-6 hours of training with feedback. The results so far are at least in qualitative agreement with predictions based on sensory bias, under which the perception of the first tone in each pair is a weighted sum of its actual frequency and its expected frequency based on previous tones [3]. However, the results could also be explained in terms of a possible response (rather than sensory) bias, based on the task that listeners were engaged in. The second experiment addresses this possibility by using similar stimuli, but with a different procedure, where the listeners’ task is to judge whether two tones had the same or different frequencies (same-different task).

### NO BIAS REVEALED WITH SAME-DIFFERENT EXPERIMENT

In this experiment, listeners were again presented with a sequential pair of tones, selected from the same range of CFs as in the first experiment. However, in this case, listeners were not asked to judge the direction of the frequency change between the tones but were simply asked whether the two tones had the same or different frequencies. In one third of trials, the two tones had the same frequency; in remaining trials, the tone frequencies were different. Of the trials with different tones, half had the higher tone first and half had the lower tone first, as in the original experiment. A sensory-bias model would predict that the low-high and high-low pairs of tones would result in the same pattern of responses that was observed in the up-down task. That is, when the tone pair is above the mean of the frequency range, the perceived frequency of the first tone will be biased downward, producing more “different” responses for Low-High trials and fewer “different” responses for High-Low trials. When the tone pair is below the mean of the frequency range, the opposite phenomenon should occur. In other words, the sensory-bias model would predict that the contraction bias should be present in a same-different task just as in the up-down task.

The mean results from the same-different task are shown in Fig. 3. An analysis of deviance revealed significant main effects of Δ*f* (χ_1_^2^ = 1325.89, *p* < 0.001) and tone order (low-high, high-low, or same) (χ_1_^2^ = 153.00, *p* < 0.001) but not CF (χ_1_^2^
= 0.65, *p* = 1. 00). The only significant interaction was between Δ*f* and tone order (χ_6_^2^
= 190.38, *p* < 0.001); all other two-and three-way interactions were not significant.

**FIGURE 3.**
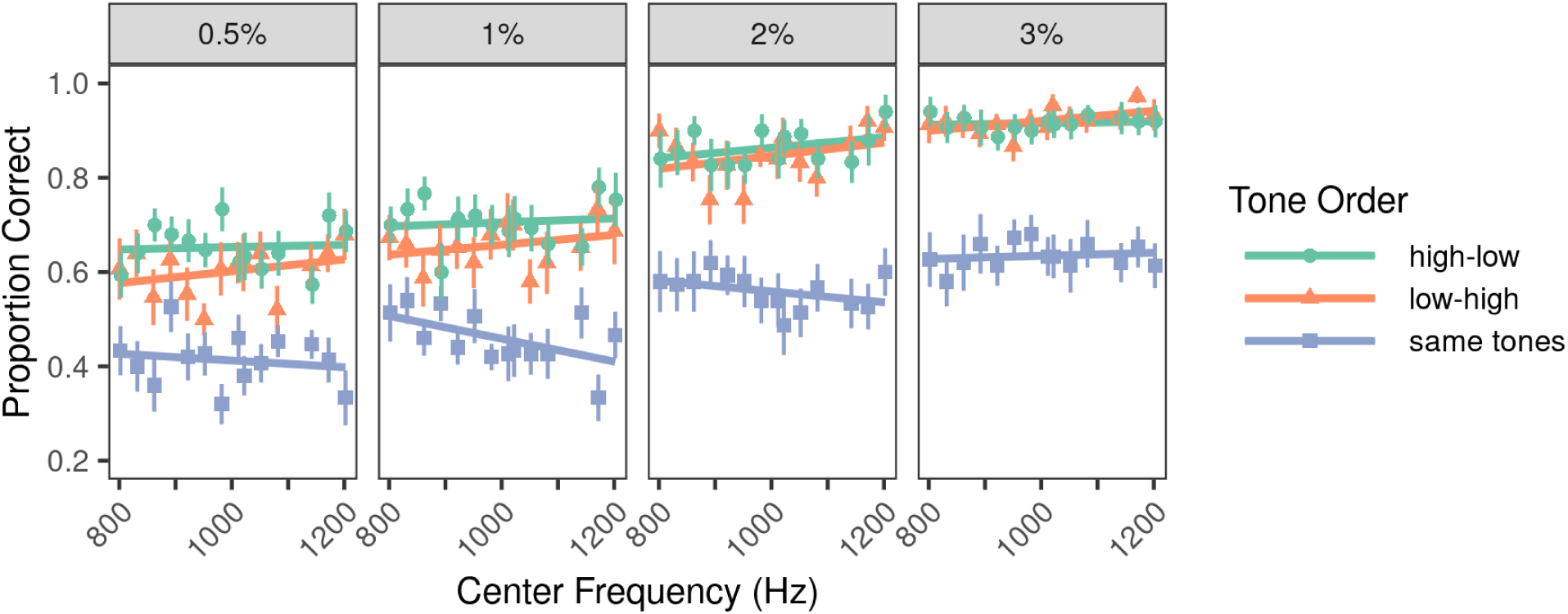
Proportion correct as a function of the center (average) frequency of the tone pair in the same-different task (N = 15). Each panel shows the results from a different value of Δ*f*. Lines represent linear regression of percent correct vs. center frequency for each tone order.

Next, linear contrast tests were performed for each Δ*f* and tone order to determine whether CF slopes were different than zero. A sensory account predicts a positive slope for the Low-High tone pairs and a negative slope for the High-Low tone pairs, as in the up-down experiment. As shown in Fig. 3, the results do not match these predictions for the Low-High or High-Low tone pairs; indeed, there seems to be no systematic effect of CF in any of the conditions. No contrast tests of CF slope in any condition reached statistical significance, suggesting that the probability of a correct response within a given tone order type did not depend on CF (all χ_1_^2^ < 2.71, all *p* = 1.00 after correction). This was also true when data were averaged across Δ*f* for High-Low (χ_1_^2^=0.89, *p* = 1. 00), Low-High (χ_1_^2^ = 2.26, *p* = 1. 00), and same-tone (χ_1_^2^ = 0.65, *p* = 1. 00) conditions.

In summary, no systematic bias was observed when measuring frequency discrimination using the same-different task. This result suggests that the original bias results reported by Raviv et al. [3] and replicated in our up-down task may reflect a response bias that depends on experimental task, rather than a sensory bias.

### MODEL PREDICTIONS BASED ON SENSORY OR RESPONSE BIAS

The data from the up-down and the same-different experiments were jointly modeled using two different computational models. The first model, based on sensory bias, instantiated the hypothesis that the estimate of the first tone frequency was biased toward the perceived mean of the frequency range (equivalently, the expected value of the directly preceding tone pairs). The second model, based on response bias or criterion shift, instantiated the hypothesis that listeners’ response criterion (and thus their tendency to respond “high-low” or “low-high”) shifted as a function of CF.

Predictions for the models of sensory bias (dashed lines) and criterion shift (solid lines) appear in Figure 4 along with the data from the up-down experiment, collapsed across task and test session. In general, the predictions of the criterion shift and sensory-bias models are nearly indistinguishable and successfully capture several key trends in the data. First, the predicted proportion correct depends on the tone pair type (low-high or high-low). Performance improves with increasing CF for low-high tone pairs and degrades with increasing CF for high-low tone pairs. According to the sensory-bias model, this is due to a shift in the estimate of the first tone frequency toward the mean of the rove range. In the criterion-shift model, the bias reflects shifts in a listener’s willingness to label a pitch shift between tones as upward or downward, depending on where in the overall frequency range the current tone pair lies. Second, the magnitude of the bias depends on Δ*f*; performance depends less on CF as Δ*f* increases. In the sensory-bias model this is because a biased estimate of the first tone frequency is less consequential when the frequency difference between the two tones is greater. In the criterion-shift model a similar phenomenon takes place; the sensory information increasingly outweighs the shifted criterion as the size of Δ*f* increases. Interestingly, both models fail to predict the continued existence of bias at the largest tested Δ*f* of 5% and also underestimate the observed bias at some intermediate values of Δ*f*, such as 1% and 2%.

**FIGURE 4.**
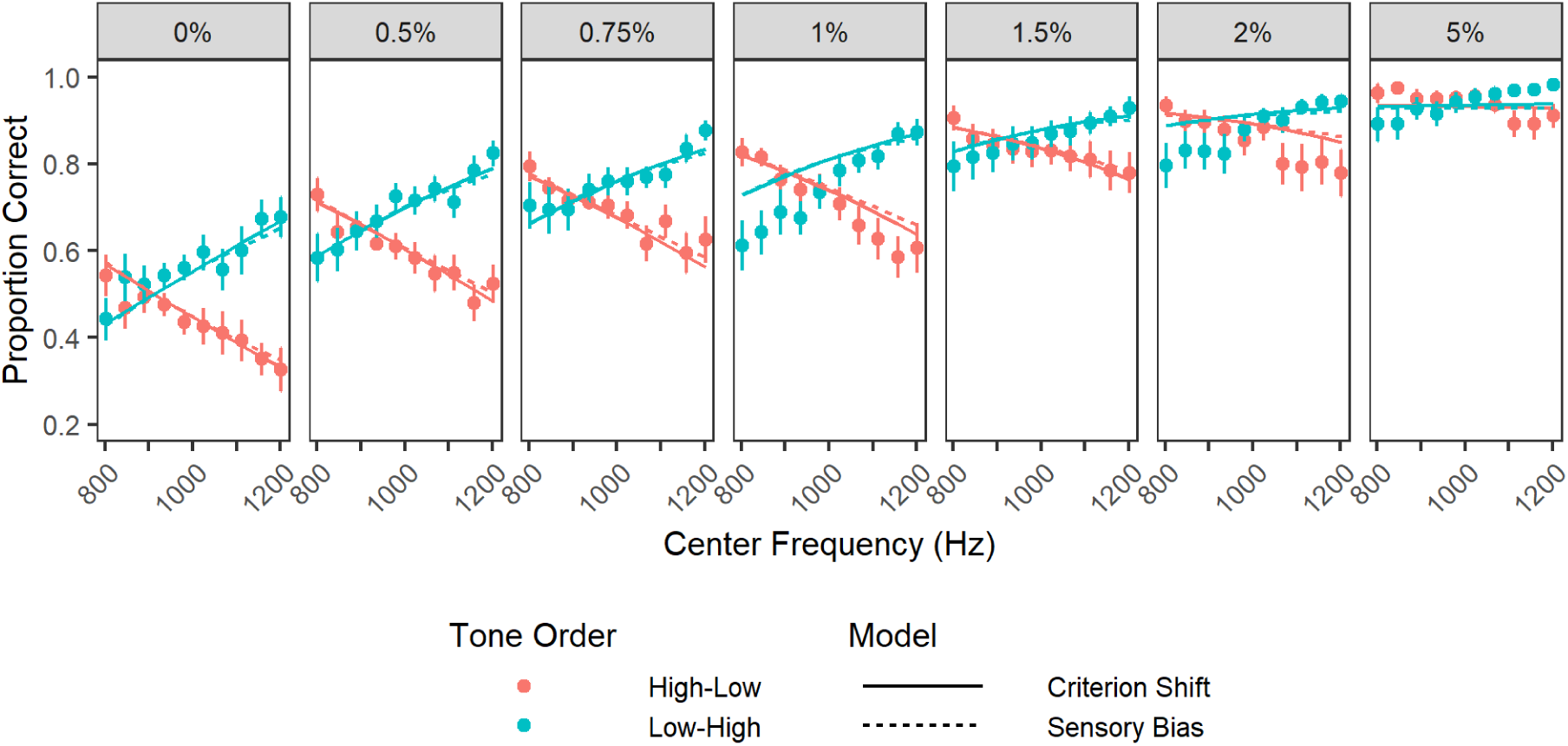
Proportion correct in the behavioral data (filled circles) and the fitted models (lines) for the up-down discrimination experiment. Each panel shows results from a different value of Δ*f*, as indicated in the grey headers. Error bars show ±1 standard error of the mean across the participants (0% Δ*f*, N = 14; 0.5% Δ*f*, N = 19; 0.75% Δ*f*, N = 19; otherwise N = 20).

Predictions of performance in the same-different experiment are shown in Figure 5 for the same two models. Both models can correctly predict that proportion of correct responses in the same-frequency trials is lower than that in different-frequency trials, and that performance improves as Δ*f* increases. The model predictions diverge, however, in terms of dependence on CF. In the sensory-bias model, the bias predicted in the up-down experiment persists in the same-different experiment because the biased estimate of the first tone frequency is made before any decision takes place. In contrast, the criterion-shift model predicts no contraction bias, consistent with the data, because changes in CF alter the model’s relative probability of responding “low-high” or “high-low” but do not alter the probability of “same” or “different” responses.

**FIGURE 5.**
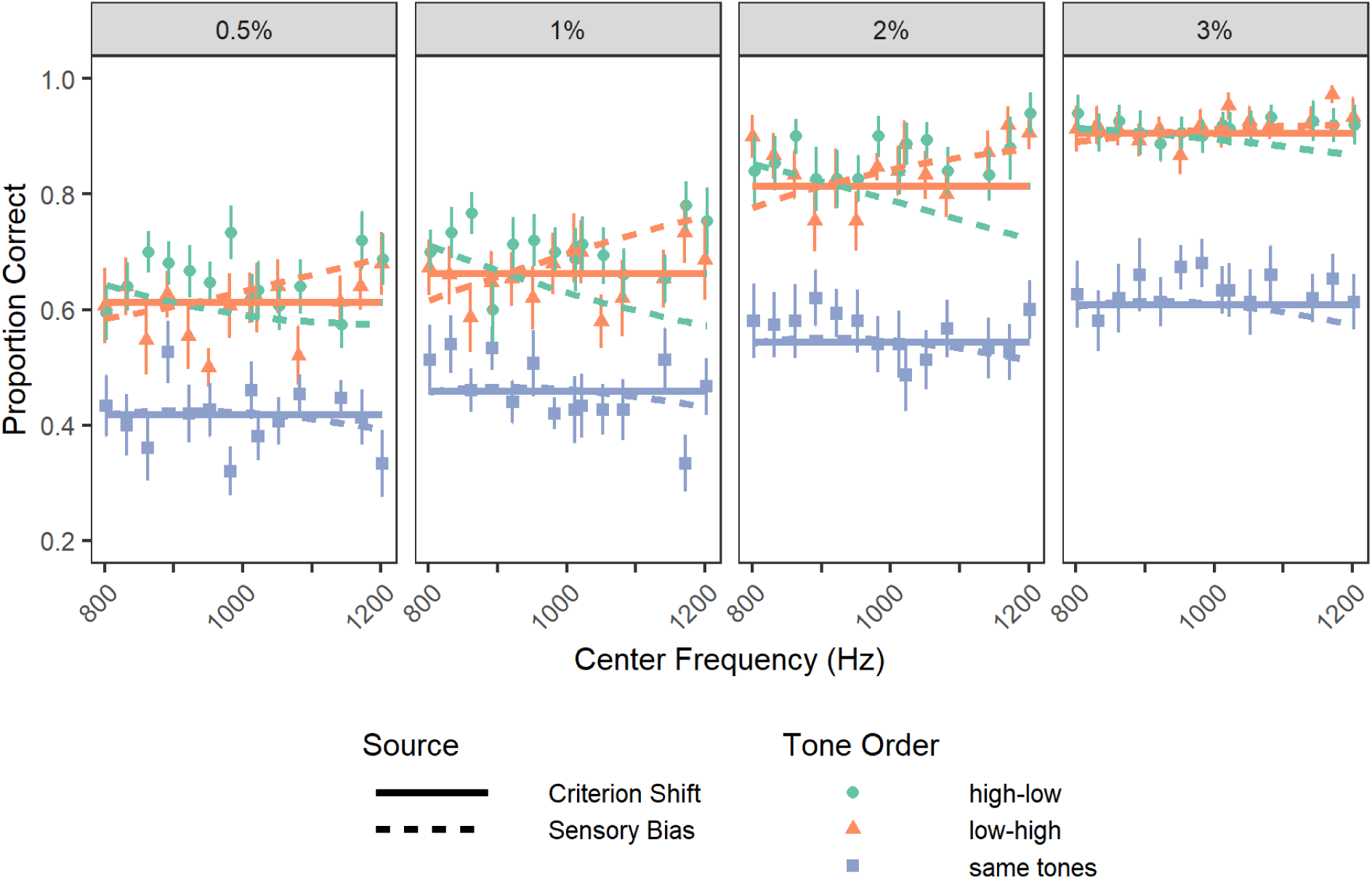
Proportion correct in the behavioral data (filled circles) and the fitted models (lines) for the same-different discrimination task. Error bars show ±1 standard error of the mean across the 15 participants.

### MODEL COMPARISONS

To provide a quantitative comparison of the sensory-bias and criterion-shift models, the data were bootstrapped 1000 times and both models were fitted to each resampled dataset. Specifically, on each repetition of the bootstrap procedure, the data were resampled at the level of individual trials with replacement and then both models were fitted to resampled data as described in the Methods. The leftmost column of Figure 6 shows histograms of the difference in the sum of squared errors between the two models separately for each task (the up-down and same-different experiments in the upper and lower panels, respectively), with positive values indicating better predictions (lower sum of squared errors) by the criterion-shift model. The criterion-shift model significantly outperforms the sensory-bias model in the up-down experiment (5% quantile = 0.05, 95% quantile = 0.09), as well as in the same-different experiment (5% quantile = 0.13, 95% quantile = 0.23). Indeed, in none of the 1000 bootstrapped samples did the sensory-bias model outperform the criterion-shift model for either task. The remaining columns of Figure 6 show histograms of the internal model parameters that were shared in both models (the “perceived mean”, “sensitivity”, and “attention” parameters; see Table 1) and demonstrate that the distributions of these parameters were similar across the two models. Both models predicted a perceived mean frequency approximately 90 Hz below the true mean of the frequency range at 1000 Hz, consistent with the “midpoint” of the contraction bias seen in Figure 2 near 900 Hz. Likewise, the estimated underlying sensitivity of the two models were similar, deviating by less than 0.05 percentage points and clustered near 0.7-0.8%, comparable to the JNDs estimated from the behavioral data shown in Figure 1. The estimated influence of sporadic inattention (or lapse rate) was similar, deviating by less than 0.02 between the two models and clustering around a proportion of 0.13-0.14, implying listeners responded randomly in 13-14% of trials. The parameters unique to each model are not visualized in Figure 5 but demonstrated similarly tight estimate distributions. Collectively, these results suggest that the amount of data available and the model fitting procedure used yielded highly stable parameter estimates and consistency between the two models in terms of parameters that were shared across the models.

**FIGURE 6.**
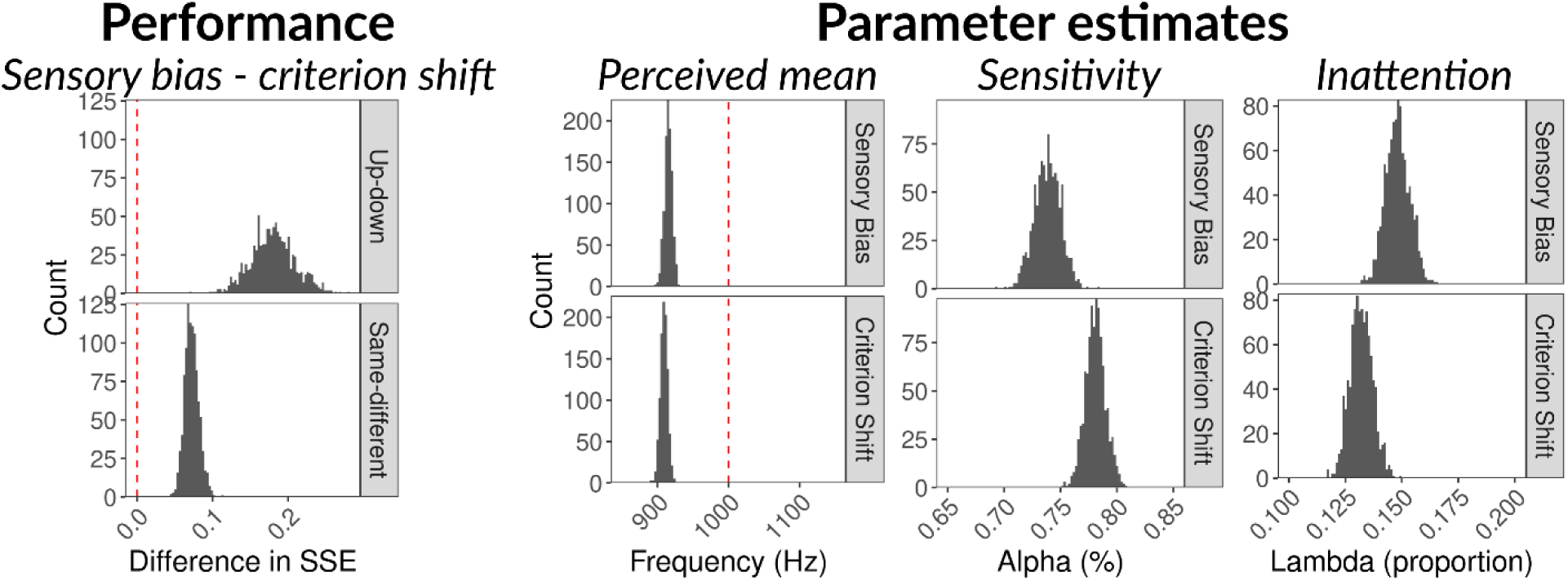
Histograms of bootstrap estimates of performance and of the three parameters shared by both the sensory bias and criterion-shift models. The difference in performance between the two models is summarized in the left-most panels as the difference of the sum of squared errors in the criterion-shift model and in the sensory-bias models, with positive values favoring the criterion-shift model. The perceived mean parameter, corresponding to *f*_mean_ in each model, was transformed from the log_10_ space in which it was tracked in the optimization procedure before visualization. The sensitivity parameter, corresponding to *α* in each model, was likewise transformed into units of percent change in frequency (Δ*f*) for visualization rather than the units in which it was tracked in the optimization procedure. The right-most panels show estimates of λ, which represents the inattention factor, or the proportion of trials for which responses were random.

**TABLE 1.** Free parameter symbol, description, and unit as well as the upper and lower bound of the range (in the transform space) that these parameters could take in the optimization procedure.

| Parameter | Short Name | Description | Transform | Lower Bound | Upper Bound |
| --- | --- | --- | --- | --- | --- |
| $\alpha$ | Sensitivity | Underlying frequency difference limen of the listener | $\log_{10}$ | -5 | 1 |
| $\lambda$ | Inattention | Proportion of time listener was not attending to task | None | 0 | 1 |
| $\omega$ | Bias magnitude | Contribution of $f_{\text{mean}}$ to the listener's estimate of $f_1$ in the sensory-bias model | None | 0 | 1 |
| $s$ | Criterion shift slope | The slope of the criterion shift with respect to distance from the tone pair to $f_{\text{mean}}$ in the criterion-shift model | None | -0.5 | 0.5 |
| $f_{\text{mean}}$ | Perceived mean | The perceived mean of the frequency range | $\log_{10}$ | 2 | 4 |

## DISCUSSION

In the present study, we replicated the key finding of Raviv et al. [3] of contraction bias in a simple up-down pure-tone frequency-discrimination experiment and extended the findings by showing that the bias persists even in listeners whose overall sensitivity is very high (an order of magnitude greater than in the Raviv et al. study) and after extensive perceptual training with feedback. However, we did not find the same bias or context effects when listeners were instructed to indicate whether the tones had either the same or different frequencies, rather than to indicate which tone had the higher or lower frequency. The fact that the contraction bias was contingent on the nature of the response being made, rather than solely on the stimuli, calls into question whether the phenomenon is truly a sensory bias, as proposed by Raviv et al. [3]. Instead, we introduced an alternative model that explained the effect in terms of a response bias and demonstrated that it provides a quantitatively better fit to the data from both experiments.

Strong psychophysical [17–21] and physiological support for sensory changes [22, 23] exist in the literature and other response-bias models (e.g., a listener consistently responds “high” or “low” when undecided) have failed in the past [24]. Hence, our present result alone do not provide conclusive evidence in favor of the criterion-shift model. However, the sensory-bias model suffers from weaknesses that must be taken into consideration. Most notably, it is unclear *a priori* why a sensory system would make a biased estimate of the frequency of the first tone given the short interstimulus intervals used here. Clément et al. [25] measured roved frequency and intensity discrimination thresholds for 50-ms pure tones and varied the silent interstimulus interval between blocks of trials from 0.5 to 10 seconds. They found that sensitivity measured in *d’* decreased from 2 (exceeding JND) to a value of 1 (equal to the JND) after 2 seconds for intensity discrimination but not until 10 seconds for frequency discrimination. Similar results have been reported in earlier studies showing that the duration of the interstimulus interval has a relatively small effect on performance in frequency discrimination task, at least for listeners without congenital amusia or cortical lesions [11, 26]. These data suggest that the memory trace for both frequencies in our experiment (950 ms ISI), which was identical to the Raviv et al. [3] experiment, should have been largely intact. In fact, a recent study found that listeners appear to guess *less* often in this task when the ISI increases from 0.5 to 2 seconds [27].

Other evidence provides indirect support for a response-bias account of frequency-discrimination context effects. For example, several studies have shown that some listeners are highly sensitive to changes in frequency but struggle to label the direction of a frequency change. This deficit is especially apparent in animals and humans with unilateral lesions of the auditory cortex, auditory midbrain, or dorsolateral frontal cortex [28], but also occurs in listeners with apparently normal auditory functioning [11, 12, 29, 30]. Moreover, two-alternative forced-choice paradigms that include a third “undecided” option, or ternary synchronous judgement, have been shown to produce more accurate estimates of sensory discrimination thresholds [31]. This suggests that at least some amount of the bias in two-alternative forced-choice tasks is under conscious control of the participant [32]. These results point to an important role of higher-level decision-making processes in pitch discrimination and suggest that context effects could emerge at higher, non-sensory, stages of processing.

In conclusion, the contraction bias appears to be a robust phenomenon in pitch discrimination tasks that persists despite high sensitivity and extensive training. However, the bias found here seems to be better understood as a response bias, where a listener’s response criterion shifts as a function of the tone frequencies, than as a sensory bias, where a listener’s estimate of the first tone frequency is influenced by the (weighted) mean of the preceding tones. These findings imply that the contraction bias measured here may reflect the influence of decision criteria rather than sensory shifts in the perception of frequency.

## MATERIALS AND METHODS

### LISTENERS

#### UP-DOWN EXPERIMENT

A total of 20 normal-hearing listeners (14 females and 6 males; 20-61 years old) participated. All listeners had normal hearing, defined as pure-tone thresholds of 20 dB HL or less at audiometric (octave) frequencies between 250 and 8,000 Hz. Listeners ranged in musical experience or training from 0-40 years (mean = 8.05 years). All 20 listeners participated in the first three test sessions. A subset of 14 (8 females, 6 males; 20-61 years; mean musical experience of 8.00 years) also completed the fourth test session. All listeners in this study provided written informed consent and were compensated for their time at an hourly wage. The protocol was approved by the Institutional Review Board of the University of Minnesota.

### SAME-DIFFERENT EXPERIMENT

Fifteen of the listeners (10 females, 5 males; 20-61 years) who had participated in the up-down experiment also participated in the same-different experiment. These listeners also ranged in musical experience or training from 0-40 years (mean = 8.33 years).

### APPARATUS

The stimuli were generated in Matlab (The Mathworks, Natick, MA), converted via a 24-bit Lynx 22 digital-to-analog converter (Lynx Studio Technology, Costa Mesa, CA) and presented diotically to listeners via HD 650 headphones (Sennheiser USA, Old Lyme, CT). All testing took place within a double-walled sound-attenuating room.

### PROCEDURE

#### UP-DOWN EXPERIMENT

In this experiment, listeners heard two pure tones that were separated by a silent temporal gap. Half of the listeners were asked to select the higher-frequency tone in the first half of the experiment and the lower-frequency tone in the second half of the experiment; the other half of the listeners performed the tasks in the opposite order. Immediately after the second tone was played, the text ‘Which tone was higher?’ or ‘Which tone was lower?’ appeared on the computer screen, depending on the task. Listeners responded by pressing “1” or “2” on a computer keyboard or clicking one of two on-screen virtual buttons using a computer mouse with no time constraint. “1” and “2” corresponded to the tone in the first and second interval, respectively. Feedback was provided (“correct” or “wrong”) immediately after each response. The next trial began 700 ms after the listener responded to the previous trial.

Stimuli in this experiment were designed to be nearly identical to those in Raviv et al. [3]. Each trial consisted of two pure tones; each pure tone was shaped by 10-ms raised-cosine onset and offset ramps had a total duration of 70 ms. The two tones were each presented at 65 dB SPL and were separated in time by 950 ms of silence. Within a block, the average frequency of the tone pair was roved between 800 and 1200 Hz. Specifically, the pure tones were centered around ten different frequencies: 800, 844, 889, 934, 978, 1022, 1066, 1110, 1156, or 1200. The tone order (Low-High or High-Low) was selected on each trial with equal *a priori* probability. The frequency difference between the tones (as a percentage of the average frequency, or CF, of the tone pair) and the target tone (“select the higher tone” or “select the lower tone”) were fixed throughout each block. Three frequency differences (Δ*f*) were tested in the first session: 1, 2, and 5%.

The first session took a total of 30 minutes, during which each listener completed six blocks of trials (3 *Δf*s x 2 task types). All Δ*fs* were tested before switching the task type from high to low and vice versa. The order of Δ*f* blocks for each target tone was randomly and independently determined at the beginning of the session for each participant. Each block was intended to contain 40 trials (10 center frequencies x 2 tone orders x 2 repetitions of each tone order at each CF), for a total of 240 trials. However, due to an experimenter error, not all participants were tested on the 889 Hz condition, and so data from the 889 Hz-condition from the first session were discarded for all participants and not analyzed.

The second, third, and fourth sessions of the up-down discrimination experiment were identical to the first session, except the center frequencies tested included 889 Hz, as originally intended, and the Δ*f*s tested differed. Each session was completed on a separate day.

For the second session, values of Δ*f* of 1, 2, and 5% were tested. Listeners completed 30 blocks (5 per *Δf* x 2 task types). Each block comprised 40 trials (10 CFs x 2 tone orders x 2 repetitions of each tone order at each CF) for a total of 1200 trials. For the third session, Δ*f*s of 0.5, 0.75, and 1.5% were tested. Listeners completed 48 blocks (8 per Δ*f* x 2 task types). Each block comprised of 40 trials (10 CFs x 2 tone orders x 2 repetitions of each tone order at each CF) for a total of 1920 trials. For the fourth and final session, Δ*f*s of 0, 0.75, and 1.5% were tested. In the 0% frequency difference condition, the first or second interval was arbitrarily assigned as the “target” with a probability of 0.5. An answer was deemed “correct” if the listener’s response matched the arbitrary assignment of the computer. If the computer arbitrarily assigned the first interval as the target interval, a “correct” message would appear on the computer monitor screen if the listener responded that the first interval contained the target. In contrast, an “incorrect” message would appear on the computer monitor if the listener indicated that the target was in the second interval. Listeners completed 48 blocks (8 per *Δf* x 2 task types). Each block comprised of 40 trials (10 CFs x 2 tone orders x 2 repetitions of each tone order at each CF) for a total of 1920 trials.

#### SAME-DIFFERENT EXPERIMENT

The same-different experiment was completed after the up-down experiment. Listeners heard two pure tones in succession. Immediately after the second tone ceased, the text ‘Were the tones the (1) same or (2) different?’ appeared on the screen. Listeners responded by pressing “1” or “2” on a computer keyboard or clicking one of two on-screen virtual buttons using a computer mouse with no time constraint. After each response, feedback was provided in the form of a message (“correct” or “wrong”) that appeared on the computer screen. The next trial commenced 700 ms after a listener’s response.

The stimuli in the same-different experiment were very similar to those used in the discrimination experiments. Each trial consisted of two pure tones, each shaped by 10-ms raised-cosine onset and offset ramps with a total duration of 70 ms. The pure tones within each pair were separated by a 950-ms silent interval. The tones were presented at 65 dB SPL and centered around one of the following frequencies (in Hz): 800, 830, 860, 890, 920, 950, 980, 1010 (another typing error, as this was supposed to be 1110), 1020, 1050, 1080, 1140, 1170, and 1200. Three types of tone pairs were tested in the same-different task: High-Low, Low-High, or same-tone (in which the frequencies of the first and second tones were both equal to the CF). Each tone order was presented twice at each CF within a block. Within a block, one Δ*f* value was tested with the Δ*f* equal to 0.5, 1, 2 or 3% of the CF. Each Δ*f* was tested in five separate blocks. The presentation order of Δ*f* was randomly determined at the beginning of the experiment for each listener. Listeners completed 20 blocks (4 Δ*f*s x 5 repetitions per Δ*f*), each of which contained 84 trials (14 CF x 3 tone orders x 2 repetitions of each tone order at each CF), for a total of 1680 trials. Listeners completed this experiment in an average of 90 minutes.

### DATA ANALYSIS AND PARAMETER ESTIMATION

### JND ANALYSIS

The just-noticeable difference (JND) refers to the threshold at which a listener can just detect a frequency difference between successive tones. JNDs were estimated using Equation 1 from Dai and Micheyl [16]. According to this equation, the proportion of correct trials (PC) can be expressed as:

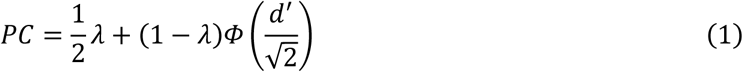

where λ is the proportion of trials during which the listener does not attend to the task (i.e., inattention or lapse rate), *dd*′ is the underlying sensitivity of the listener, and Φ is the normal cumulative distribution function. According to this equation, performance is fixed at chance during inattention. The listener’s sensitivity is assumed to be proportional to the log-transformed difference in the frequencies of the two tones:

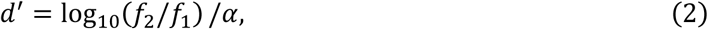

where *α* is a free parameter and *f*_1_ and *f*_2_ are the frequencies of the first and second tones, respectively. A few points must be made here. First, note that Equation 1 implicitly assumes that *f*_2_ > *f*_1_. For trials where *f*_2_ < *f*_1_, we invert the sign of *d*′ before evaluating Equation 1. Second, at least over the range of frequencies in question, log_10_(*f*_2_/*f*_1_) is approximately proportional to the Δ*f*/*f* used in Dai and Micheyl [16]. Finally, as is standard in the analysis of frequency discrimination data [16], we assume that the sensory estimates of *f*_2_ and *f*_1_ are normally distributed and have equal variance.

The curves and JND estimates that appear in Figure 1 were generated by dividing the entire dataset into subsets according to test session and then estimating a JND for each subset by optimizing the parameters in Equation 1 to minimize the sum of squared errors between the output of the equation and the observed data. This optimization procedure was performed in the R programming language [33] using a bounded Hooke-Jeeves optimizer implemented via the *pracma* package [34].

To test whether the JND changed as a function of test session and target tone, all data was analyzed according to subject, test session, and target tone, and this process was repeated to estimate the JND for each subset. The resulting JND estimates were then used as the response variable in a linear mixed-effects model with fixed effects of test session and target tone and a random intercept for each listener. The linear mixed-effects model was fit in R 3.5.1 using the *lme4* package [35]. The model was fitted via penalized maximum likelihood estimation. The model was analyzed in two ways. First, to determine the significance of the fixed effects, we calculated *F*-tests in a Type II analysis of deviance using the *Anova* function provided by the *car* package [36]. The degrees of freedom for the *F*-tests were selected using the Kenward-Rogers method implemented in the *pbkrtest* package [37]. Second, to assess the significance of linear contrasts between conditions, we calculated *F*-tests via the *phia* package [38], again using the Kenward-Rogers method for selecting the degrees of freedom. Finally, all p-values associated with each model were jointly corrected using the Holm-Bonferroni method [39]. Corrected p-values were compared against a criterion of 0.05 to determine statistical significance.

### PERCENT CORRECT ANALYSIS

To analyze the data from the discrimination experiments in terms of how percent correct varied as a function of the experimental parameters, generalized linear mixed-effects models were fitted to the percent correct data. These models were fitted and analyzed in the same way as the mixed-effects model used to analyze the JND estimates, with the following exceptions. First, because the data were dichotomous in nature (i.e., each trial was correct or incorrect), we used a logistic link function. Second, because the Kenward-Rogers method for selecting degrees of freedom is unavailable in R for generalized linear-mixed effects models, Wald χ^2^-tests were used instead of *F*-tests.

In the model fit to the up-down discrimination data, fixed effects included center frequency (CF), tone order, target tone, and Δ*f*, while random effects included intercepts, tone order, and the interaction between CF and tone order for each subject. Notably, we collapsed data across sessions for this analysis. In the model fit to the same-different data, fixed effects included Δ*f*, trial type (same, High-Low, or Low-High), and CF with random effects of intercepts and the interaction between trial type and CF for each subject.

### COMPUTATIONAL MODELS

Each model shared several features. First, the models were fit to all data, i.e., the individual trial-level data for every subject and condition. The outcome variable was binary in nature; a trial was either correct or incorrect. Second, free parameters were optimized *jointly* for the up-down and same-different discrimination experiments by minimizing the sum of squared errors between the model output and the observed data. This optimization procedure was performed in the R programming language [33] using a bounded Hooke-Jeeves optimizer implemented via the *pracma* package [34].

### SENSORY-BIAS MODEL

#### UP-DOWN DISCRIMINATION

In the sensory-bias model, it was assumed that listeners’ estimates of the frequency of the first tone (*f*_1_) were biased toward the perceived mean of the frequency range of the tones. The proportion of correct trials in the sensory-bias model of the up-down task was:

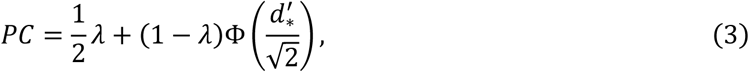

where *λ* is the inattention factor, or the proportion of trials for which listeners are responding randomly, and *d* indicates the sensitivity of the listener. *d*^′^ is defined as:

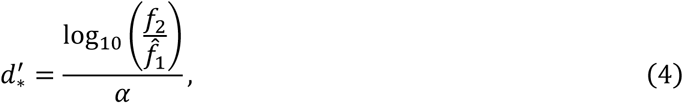

where *α* represents the frequency JND in units corresponding to those of the numerator, *f*_2_ indicates the frequency of the second tone, and 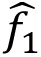 indicates the (potentially biased) estimate of the frequency of the first tone. As in Equation 1, we invert the sign of d’_*_ if *f*_2_< *f*_1_ Note that the only difference between the simple PC equation (Equation 1) and present PC equation is the replacement of *d*′ with *d*^′^. Then:

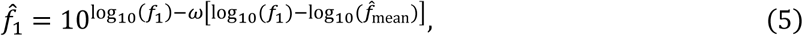

where *f*_1_ corresponds to the true frequency of the first tone and *f*^_mean_ corresponds to the perceived mean of the frequency range. In effect, the sensory-bias model assumes that listeners use an estimate of *f*_1_ that is biased toward the perceived mean *f*^_mean_ to an extent controlled by parameter *ω*. If *ω* = 0, Equation 3 reduces to Equation 1. In contrast, if *ω* = 1, then listeners are assumed to completely ignore sensory information about *f*_1_ and instead base their estimate of *f*_1_ exclusively on the perceived mean of the frequency range.

Parameters that could vary while fitting Equation 3 appear in Table 1. Additionally, any transformations applied to parameters before fitting and the range of values free parameters were allowed to take are indicated in Table 1. Parameters used only in other models (described below) are also shown in Table 1.

### SAME-DIFFERENT DISCRIMINATION

Performance in the same-different experiment was modeled using a differencing strategy based on signal detection theory [40, 41]. Specifically, the estimated frequency of one tone was subtracted from the estimated frequency of the other and then the difference was compared to a criterion. Stimuli were judged as “different” if the absolute difference exceeded the criterion; otherwise, stimuli were judged to be the “same.” Therefore, overall performance in the same-different task was modeled as:

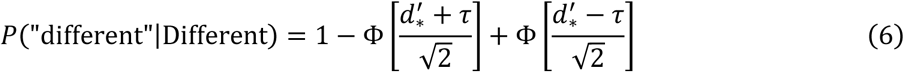

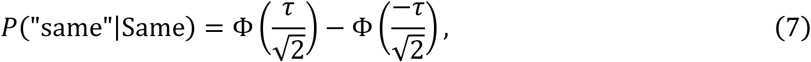

where *τ* represents the decision criterion, “different” and “same” represent responses of different and same, respectively, and Different and Same represent different-frequency and same-frequency trials, respectively. Note that the same *d*^′^ value used in Equation 3 is used here. As such, the same-different model should exhibit the same basic bias patterns as the up-down model. Values of *τ* were derived from data. One value was derived for each *Δf* tested from mean false-alarm rates in same-tone trials. These values were derived by solving Eq 9.8 in [40] for *τ* (referred to as *k* in [40]) as:

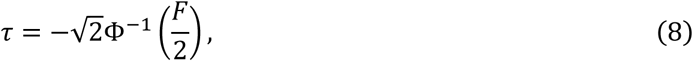

where *F* indicates the false alarm rate. Those derived values were 0.543, 0.613, 0.771, and 0.905 for 0.5%, 1.0%, 2.0%, and 3.0% trials, respectively. Since the up-down and same-different pitch-discrimination tasks were modeled jointly, the free parameters were restricted to the same ranges as indicated in Table 1.

### CRITERION-SHIFT MODEL

#### UP-DOWN DISCRIMINATION

In our signal detection theory model, the decision variable in the up-down task is the difference of sensory estimates of the two (log-transformed) tone frequencies 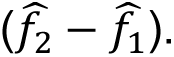 The listener’s task is to decide whether the derived decision variable on any given trial belongs to the distribution generated by Low-High trials (with a mean 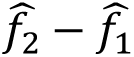 above 0) or the distribution generated by High-Low trials (with a mean 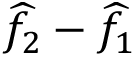 below 0). If a listener is unbiased, then their decision criterion *k* is located at zero (and in the limit they will make the same number of incorrect responses in Low-High trials and in High-Low trials). If a listener’s criterion shifts downward, they will respond “low-high” more often and their pattern of errors will change. Specifically, they will correctly identify more Low-High trials as Low-High trials (hits) but also misidentify more High-Low trials as Low-High trials (false alarms). The opposite pattern of results will occur if a listener’s criterion shifts upward.

We hypothesized that the contraction bias observed in this task was due to systematic changes in the listeners’ decision criterion, as opposed to changes in their sensory estimates of *f*_1_ and *f*_2_. That is, listeners shifted their willingness to respond “high-low” or “low-high” as a function of the position of the tone pair relative to the mean of the frequency range. Specifically, we hypothesized that listeners would tend to respond “low-high” more often for tone pairs above the mean of the frequency range and “high-low” more often for tone pairs below the mean of the frequency range. This can be justified intuitively by assuming that listeners are in part responding to the question: “Was the current tone pair higher or lower than the previous tone pair(s)?”. To this end, we modeled the listeners’ decision criterion *k* as a function of the distance between the tone pair and the perceived mean of the frequency range (Equation 9).

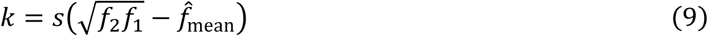

In Equation 9, a negative value of *ss*, the slope of the criterion shift, would result in listeners being more likely to respond “low-high” when the tone pair was above the mean and more likely to respond “high-low” when the tone pair was below the mean (as in the contraction bias). Proportion correct was calculated for the up-down criterion-shift model (separately for High-Low and Low-High trials) as…

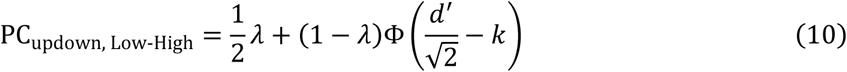

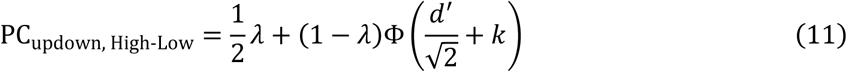

A schematic of the criterion-shift model appears in Figure 7, where shifts in biased criterion (red) are depicted as a function of the position of the tone pair in the rove range relative to the perceptual mean.

**FIGURE 7.**
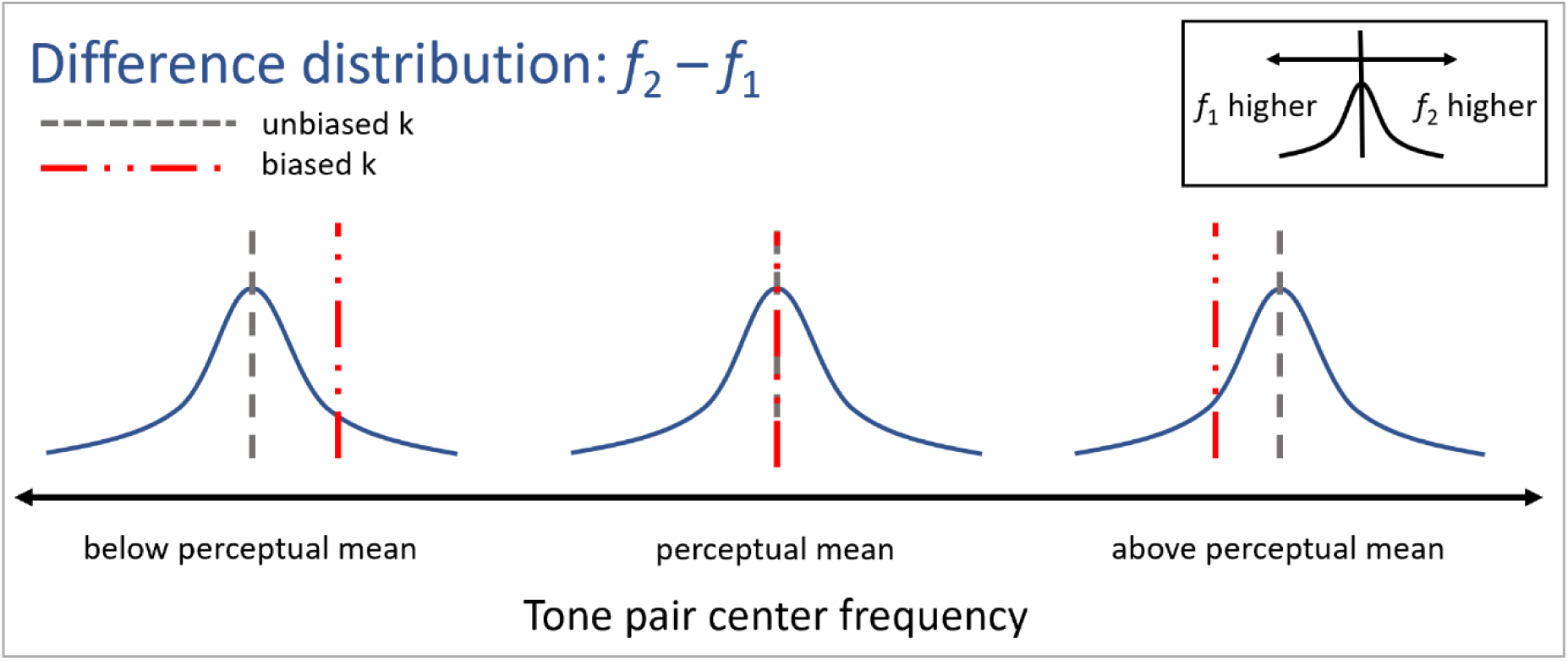
Schematic of the criterion-shift model. The black line represents the continuum of tone pair center frequencies, where the center of the line represents the perceptual mean. Blue curves depict the *f*_2_-*f*_1_ difference distribution. Gray and red vertical dashed lines correspond to unbiased and biased criterion, respectively. Area beneath the curve to the right of a line corresponds to the probability that *f*_2_ is higher than *f*_1_; area beneath the curve to the left of a line corresponds to the probability that *f*_1_ is higher than *f*_2_.

### SAME DIFFERENT EXPERIMENT

Performance in the same-different experiment under the criterion-shift model was calculated as:

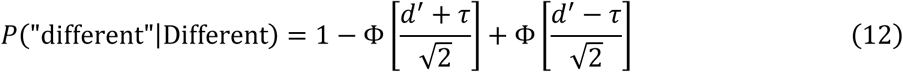

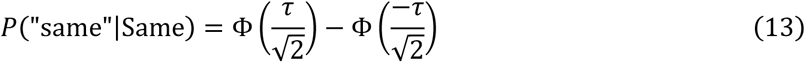

Note that these equations are the same as Equations 6 and 7 except using *d*′’ instead of d’_*_. Because there is no judgment of relative frequency involved, there is no shift in criterion associated with changes in CF. As such, Equations 12 and 13 do not demonstrate a contraction bias and predict the same results at all CFs.

